# Atomistic simulation of protein evolution reveals sequence covariation and time-dependent fluctuations of site-specific substitution rates

**DOI:** 10.1101/2022.06.01.494278

**Authors:** Christoffer Norn, Ingemar André

## Abstract

Thermodynamic stability is a crucial fitness constraint in protein evolution and is a central factor in shaping the sequence landscapes of proteins. The correlation between stability and molecular fitness depends on the mechanism that relates the biophysical property with biological function. In the simplest case, stability and fitness are related by the amount of folded protein. However, when proteins are toxic in the unfolded state, the fitness function shifts, resulting in higher stability under mutation-selection balance. Likewise, a higher population size results in a similar change in protein stability, as it magnifies the effect of the selection pressure in evolutionary dynamics. This study investigates how such factors affect the evolution of protein stability, site-specific mutation rates, and residue-residue covariation. To simulate evolutionary trajectories with realistic modeling of protein energetics, we develop an all-atom simulator of protein evolution, RosettaEvolve. By evolving proteins under different fitness functions, we can study how the fitness function affects the distribution of proposed and accepted mutations, site-specific rates, and the prevalence of correlated amino acid substitutions. We demonstrate that fitness pressure affects the proposal distribution of mutational effects, that changes in stability can largely explain variations in site-specific substitution rates in evolutionary trajectories, and that increased fitness pressure results in a stronger covariation signal. Our results give mechanistic insight into the evolutionary consequences of variation in protein stability and provide a basis to rationalize the strong covariation signal observed in natural sequence alignments.

## Introduction

Unstable proteins reduce cell fitness by either reducing the concentration of active molecules or due to toxicity of the unfolded state[1–3]. Therefore thermodynamic stability provides a link between structure and energetics in proteins on the one hand, and fitness of sequences sampled during the process of evolution on the other hand. There is ample evidence to suggest that the stability fitness constraint is an important factor controlling the evolution of protein sequences. Models based on stability fitness constraints can explain the variation in overall substitution rates between proteins[1, 4, 5], mutational rates at individual sites in proteins[6, 7], and global amino acid substitution patterns [7].

The impact of thermodynamic stability on molecular fitness depends on the mechanism that couples stability to fitness. If the unfolded state is toxic, the cytotoxicity and the concentration of the unfolded state control the fitness effect of a stability-altering mutation. The effective population size also determines the impact of the fitness effect, as fixation probability for small selection coefficients is a function of effective population size and selection coefficient. Because proteins span a wide range of toxicity and expression levels, and there is considerable diversity in effective population sizes between organisms, it is important to understand the impact of fitness pressure on protein evolution. Here we study how variations in fitness pressure impact the sequence evolution of proteins.

Previous work to understand the effect of protein stability on evolution has primarily used simplified models of protein energetics on rigid protein backbones. Given the high sensitivity of real proteins to even the most conservative mutations, it is valuable to develop atomistic models of protein evolution that capture the detailed energetics and conformational variability of proteins. Here, we develop an all-atom evolution simulator, RosettaEvolve, to simulate evolutionary trajectories using the Rosetta macromolecular modeling package[8, 9]. We use RosettaEvolve to study how fluctuations in protein stability change in site-specific mutational rates with a single fitness function. Next, we study how selection pressure affects the distribution of proposed and accepted mutational effects and the degree of residue-residue covariation observed in evolved protein sequences.

To generate evolutionary trajectories in RosettaEvolve a fitness model is used to evaluate how a change in computed stability affects the fitness of a sequence variant. The fitness model is then coupled with an evolutionary dynamics model that evaluates the fate of mutations during stochastic sequence trajectories. There are two possible fates under strong selection and weak mutation: Either the mutation spreads through the population and becomes fixed or is lost due to drift or selection. The probability of fixating a proposed mutation depends on the fitness of the new protein sequence but also on the effective population size. We evaluate fixation probabilities with Kimura’s expression for fixation probability[10]. Based on these two elements - the stability fitness model and expression for fixation probability - it is possible to describe the evolution of protein sequences and predict substitution rates. Substitution rates, amino acid substitution rates, and site-specific rates can be calculated directly from stochastic trajectories[1, 2, 4, 5, 11–18] or by converting the stability model into a Markov state model[6, 7].

We have recently developed a Markov state model to predict amino acid substitution and site-specific rates in proteins[7]. In this approach, substitution rates are evaluated in the context of a single structural and sequence environment, typically based on the native protein structure and sequence. In this study, we generate evolutionary trajectories with RosettaEvolve and use the Markov model to calculate site-specific rates along the trajectory, combining the strength of these two different approaches.

Simulation of evolutionary trajectories enables studies of how temporal and structural fluctuations affect the evolution of proteins. Prior work, which applied an approximate representation of protein structure, has shown that changes in propensities of amino acids can occur in a protein during evolution and that the protein adopts to the mutated site making reversals less favorable over time, a mechanism referred to as the evolutionary Stokes-Shift[19].

Here we show how RosettaEvolve can be used to simulate evolutionary trajectories guided by an all-atom structural model of the detailed energetics of proteins. We demonstrate how variation in the molecular fitness parameters – such as cytotoxicity, the concentration of unfolded protein, and population size - affects both the proposal and fixated distribution mutational effects. We show that site-specific mutational rates fluctuate over trajectories largely dependent on fluctuations in the stability of the protein. We also show that phylogenetic trees generated by RosetteEvolve result in a robust residue-residue covariation signal which depends on selection pressure.

## Results

### Simulation of evolutionary trajectories with RosettaEvolve

RosettaEvolve simulates evolution at the nucleotide level (Fig. 1). Differences in the chemical properties of nucleotides result in different rates for transitions and transversions [1, 2]. This bias is controlled in the simulation by specifying the transition/transversion rate ratio k. Multi-nucleotide or whole-codon changes are also observed in nature due to a multitude of genomic processes such as insertions, deletions, UV-damage, and tandem mutations[3, 4]. These nucleotide changes are captured by a multi-codon mutation rate r. The relative rate of single base pair changes to multi-codon mutation depends on r and k but also the number of states that are accessible for the multi-nucleotide route: We calculate the probability of multi-nucleotide changes as

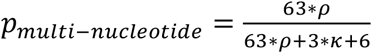

k and r are parameters in the simulation. In this study, we have set the values found to optimize the correlations with empirical amino acid substitution rates presented in Norn et. al.[5], k=2.7 and r=0.1.

**Figure 1:**
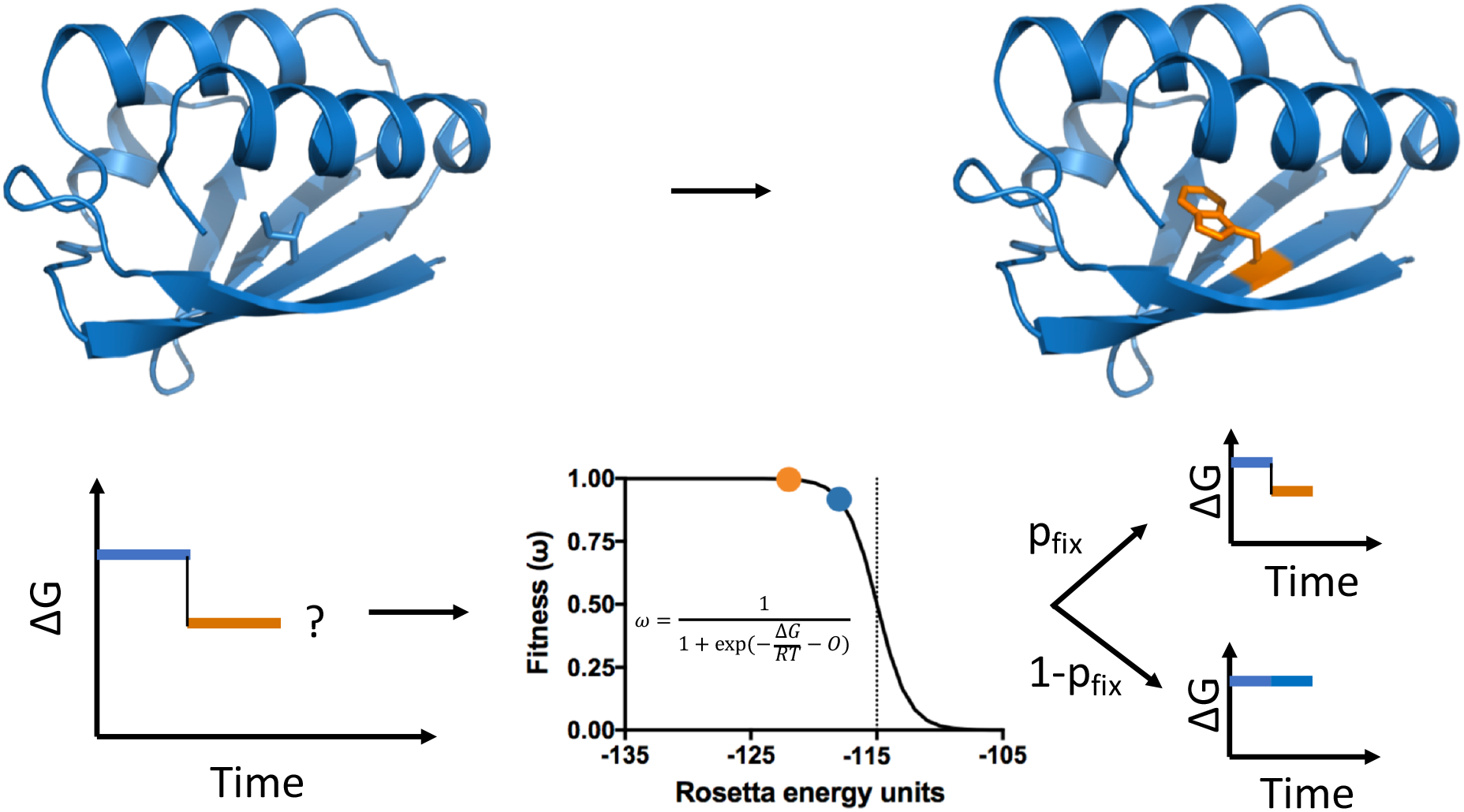
Simulation of evolutionary trajectories with RosettaEvolve. Mutations are proposed at the nucleotide level, and nucleotide changes are translated into amino acid substitutions. The fitness of a mutation is estimated by calculating the change in stability of the protein with a ΔΔG prediction method using the Rosetta all-atom energy function. Based on the change in fitness (selection coefficient), the probability of fixating the proposed nucleotide/amino acid is evaluated.

To evaluate the probability that the introduced mutation will be fixated, we first have to evaluate the fitness of the mutation. Several fitness models based on protein stabilities have been described[6–9], and RosettaEvolve can easily be extended to use alternative fitness expressions. In this study, we use a fitness model that assumes that a protein’s contribution to fitness is proportional to the fraction of the protein folded in its native conformation[9]. As described in Norn et. al.[5], for stable proteins this is mathematically equivalent to a cytotoxicity fitness model[6], where fitness depends on the concentration of unfolded protein, but with an offset ΔG. Equating fitness to the fraction folded, the expression for fitness becomes

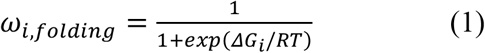

Where *ω_i,folding_* is the fitness of sequence *i* and *ΔG* = *G_folded_* – *G_unfolded_* is the free energy of folding. When *ΔG* < −3 kcal/mol, the cytotoxicity fitness model has the same mathematical form as the stability fitness function[5]

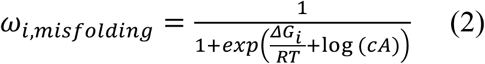

Where *c* is a toxicity parameter and *A* is the protein abundancy[6].

There are currently no methods that can accurately compute DG values with an energy function or force field. However, DDG prediction methods can reach useful correlations between computed and experimental values (r^2^=0.56 reported for the method used in this study[10]). The stability of a protein sequence after each mutation is evaluated

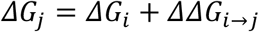

For a trajectory started from the native sequence, we must assign a stability to the native state. This is done by subtracting an offset (E_ref_) from the energy of the native sequence *E_rosetta_* so that *ΔG* = *E_rosetta_* – *E_ref_*. Analogously, as seen in equation 2, the cytotoxicity and abundance parameters offset DG in the fitness function. Furthermore, changes in effective population size (N) has a similar effect as offsetting *ΔG,* as *ΔG* ~ – log *N* [5,11]. Hence, we can model the fitness of a sequence as

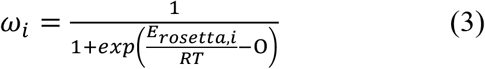

where *O* is the linear offset. Setting *O*=*E_ref_* converts the fitness function into the fraction folded model in equation 1. *O* is treated as a parameter in our simulations, while *E_rosetta_* is calculated from the structure with the Rosetta energy function. The value of *O* controls the offset of the fitness function (through the cytotoxicity/abundance parameters or effective population size) and anchors the computed energy on the free energy scale. Setting a low value of the offset assigns a low fitness to the native sequence, which forces the introduction of stabilizing mutations to increase the fitness of protein and a decrease in the energy of the protein. Conversely, setting a high value for the offset assigns a high fitness of the native sequence, which facilitates the introduction of destabilizing mutations since such mutations carry little fitness cost, leading to an increase in the energy of the protein until mutation-selection balance is reached. By sliding the value of the offset parameter, we change the effective selection pressure. From here on, we refer to the negative of the offset O as the selection pressure.

The flexibility of the Rosetta program enables different methods to calculate DDG of mutations during the evolutionary trajectory. Currently, we have implemented two methods for DDG predictions in RosettaEvolve: One with limited and one with more extensive backbone flexibility. The simulations presented in this study were carried out using the method with limited backbone flexibility, a slight variation of the DDG prediction approach in Rosetta presented by Park et. al.[10]. The method involves repacking residues that are energetically coupled to the mutated amino acid and backbone energy minimization of the focal site and the nearest neighbors in the sequence.

If populations evolve under sufficient strong selection pressure and at a sufficiently low mutation rate, the fixation probability can be estimated using Kimura’s fixation probability equation[10]. For diploid organisms

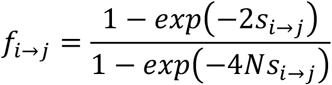

where *f*_*i*→*j*_ is the probability of fixation of a mutation *i* to *j, s*_*i*→*j*_ = *ω_j_/ω_i_* – 1 is the selection coefficient, and *N* is the effective population size. In the simulations presented here, we set *N*=10^4.2^, a value we previously found to optimize correlations between computed and empirical amino acid substitution rates [5, 11]. In experiments where we modeled changes to the selection pressure, we kept N constant and just modified the offset parameter *O*.

Computed fixation probabilities are generally too low to enable efficient simulation of evolutionary trajectories. Instead, we used a rescaled fixation probability, 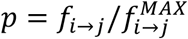, with the scale-factor defined as,

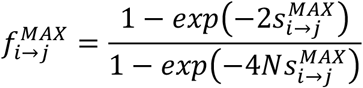

where 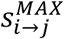 is calculated with *ω_j_* = 1, the maximal possible fitness. The scale-factor is stored to enable later rescaling for rate-calculations.

In the following sections, we apply RosettaEvolve to study a specific protein, Azurin of *P. aeroginosa.* Azurin is a 128-residue protein with an immunoglobulin-like fold. The protein has a copper-binding site and a single disulfide bond. Before generating evolutionary trajectories with azurin the protein was adapted to the Rosetta energy function with a structure refinement calculation.

### Equilibration of trajectories

Before analyzing the dynamics of sequence evolution, the simulations must be equilibrated so that the recorded trajectory is under mutation-selection balance. The fitness equilibria shift depending on the assumed stability of the protein. In our approach, the selection pressure is controlled by the offset value. A separate equilibration is required for each selection pressure. Mutations are evaluated using a ΔΔG prediction protocol that involves structure remodeling and energy minimization. This means that structural changes across the trajectory and the effect of this flexibility must be equilibrated. In principle, one could return to the starting structure after each mutation, but this requires more extensive backbone sampling and leads to a far noisier energy estimation.

To follow the progress towards equilibrium, we measure the energy, mean change in energy of accepted mutations, and the average energy rank of the amino acid selected at sites. To calculate the rank, the amino acid variants at sites are sorted according to their relative energy. The energy rank is the position in the list (1-20) for the currently selected amino acid. We ran equilibration trajectories for 11 different selection pressure values corresponding to 10 mutations/site branch length. At every integer branch length, the average change in energy relative to the best choice amino acid and the average rank was evaluated. Figure 2A shows the result for a selection pressure value corresponding to lower stability than the native sequence. Destabilizing interactions are initially introduced into the protein, which increases the mean energy rank of the current amino acid. Trajectories with varying selection pressure values will equilibrate at different protein stabilities. This is observed in Figure 2B where the average energy value for accepted sequences is plotted against selection pressure values. The mean sequence energy is linearly dependent on the selection pressure. At low selection pressure values, sequences have increased stability relative to the native sequence and the mean rank is low because the energetically best choice amino acid occurs very frequently at sites in the protein. At high selection pressure values, the mean rank is close to 10, which is the value expected with a completely random distribution of amino acids at sites. Strong selection pressures – high cytotoxicity/high abundance of the unfolded protein and/or large effective population size – thus result in proteins with increased thermal stability.

**Figure 2:**
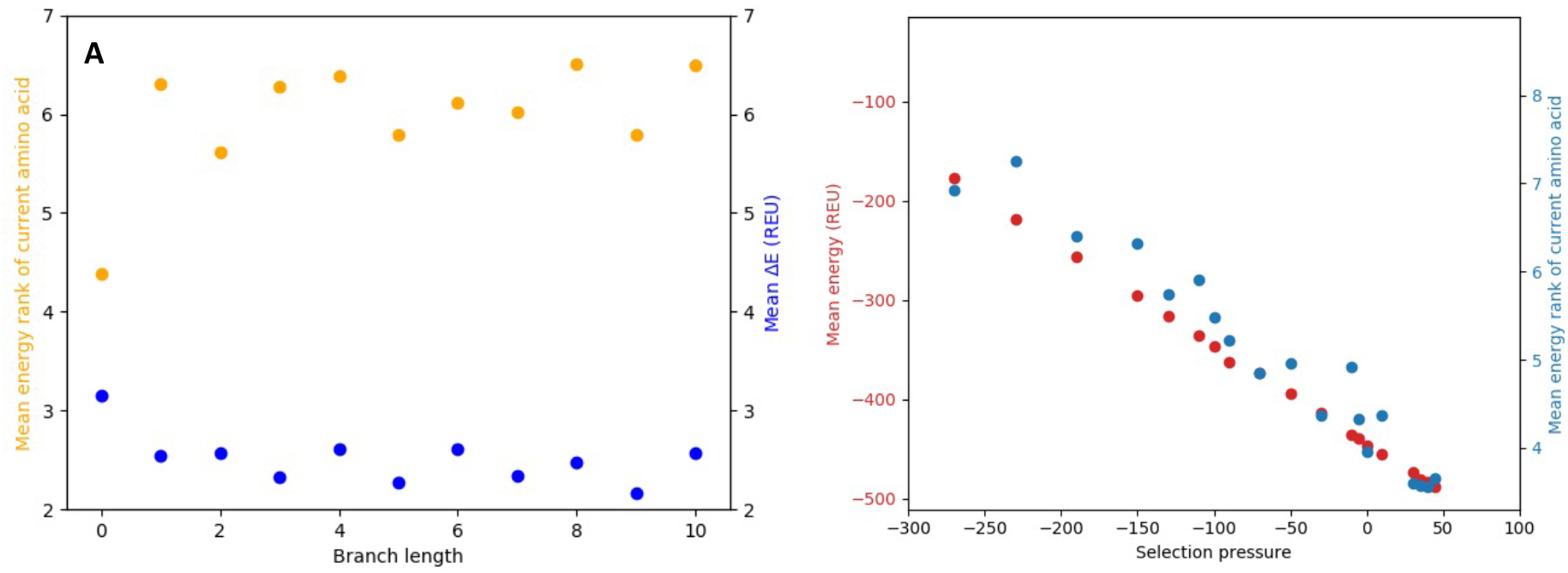
Equilibration of azurin. **A)** Mean change in energy (blue) and mean energy rank (orange) as a function of branch length for a simulation with selection pressure set to −162. **B)** Dependence of mean energy of accepted sequences (red) and mean rank (blue) on the selection pressure (fitness function offset).

### The selection pressure impacts the probability distribution over proposed and accepted DDG values

Evolutionary trajectories at different selection pressure values were generated based on the final structure at the end of the equilibration runs. From these trajectories, we summarized the probability distribution over proposed and accepted ΔΔG values as a function of the selection pressure value (Fig. 3). With decreasing selection pressure (resulting in higher protein stability), the mean energetic effect of mutation (ΔΔG) increases (Figure 3C). In other words, mutations become more detrimental when the protein stability increases. The distribution of mutational effects in real proteins behaves the same way albeit the increase of the detrimental effect is about 10 times higher [4].

**Figure 3:**
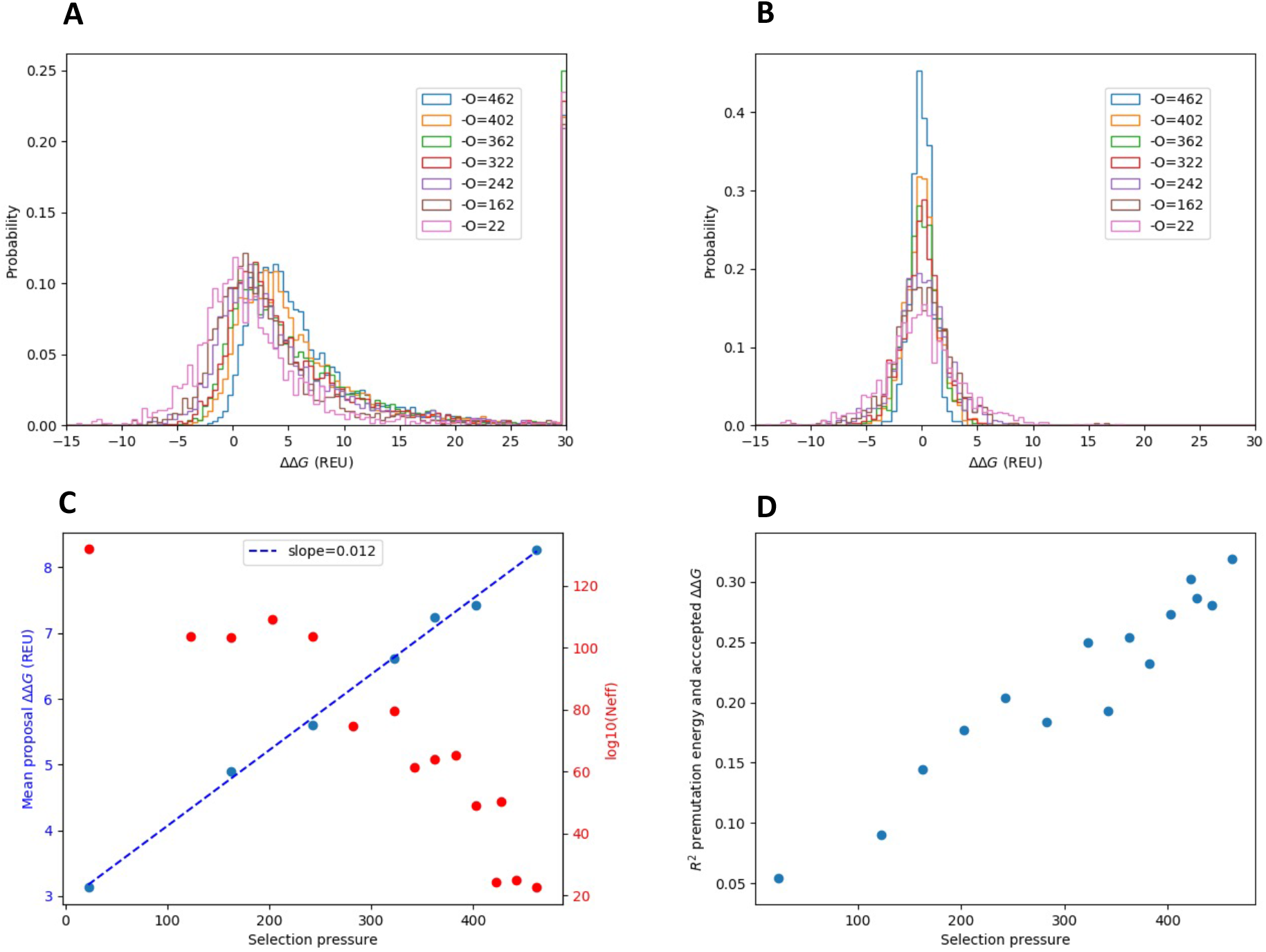
The selection pressure affects the DDG proposal and acceptance probability distribution. **A)** Proposal DDG probability distribution as function of selection pressures. Values above 100 energy units (corresponding to severe atomic clashes) were placed in the highest bin. **B)** Accepted DDG probability distribution as function of selection pressure. **C)** Mean proposal DDG value (correspond to the distribution < 100 energy units) as a function of selection pressure(blue), with a fitted line (blue). The logarithm of number of available sequences as a function of selection pressure (red) calculated with an assumption of independent sites in the protein. Energies in Rosetta Energy Units (REU). **D)** Correlation between premutation energy and accepted DDG values as a function of selection pressure. Correlations are measured as squared Pearson correlation coefficients.

At very high selection pressure values (resulting in low protein stabilities) the probability distribution over DDG for proposed mutations is symmetric and centered around 0, with an equal probability of proposing stabilizing and destabilizing mutations. Under these conditions, the distribution over proposed and accepted DDG values are almost identical. At lower selection pressures (resulting in higher protein stabilities), highly stabilizing mutations are much less likely to be proposed, and the probability distribution is shifted towards more destabilizing mutations (Fig. 3A). The probability distribution over accepted DDG values is symmetric around 0 for all selection pressure values but becomes more peaked as the stability increase (Fig. 3B).

The mean of the proposed DDG values linearly depends on selection pressure (and therefore on the mean stability of the protein, see figure 3C). Why does the proposal probability change with the stability of the protein? At high stability, there are few accessible mutations that can stabilize the protein since the best choice amino acid is often present at sites in the protein. We can estimate the space of accessible sequence at different reference stability values by multiplying together the effective number of amino acids at each site in the protein (assuming independent sites) calculated from the equilibrium amino acid frequency distribution at each site. As shown in Figure 3C, the sequence space is much smaller for more stable proteins evolved under higher selection pressure. This reduction in sequence space is likely to explain the shift of proposal DDG values towards more destabilizing mutations at higher selection pressures.

The consequence of protein stability on the probability distributions over proposed and accepted DDG-values has previously been studied by Goldstein using a contact-based energy model [12]. He found that the stability of the protein before mutation and the DDG of accepted mutations correlated. We observe the same correlation with the all-atom simulations as seen in Figure 3D: Mutations accepted in a stable protein will generally be less stabilizing than those accepted in a protein with lower stability. The correlation between premutation stability and DDG reduces with decreasing selection pressure.

### A strong covariation signal is found when phylogenetic trees are simulated by RosettaEvolve

The success of covariation analysis in identifying residue-residue contacts suggests that epistasis and coevolution are pervasive elements of evolution [25]. Yet, covariation, as measured by statistical coupling methods [26-29], is not necessarily the same as coevolution [30]. Statistical coupling methods are based on sequence alignments and do not consider that substitution has occurred along branches of phylogenetic trees. Tree-based methods to detect coevolution based on evolutionary theory have been developed [31, 32], but their high computational cost hampers their use. A method developed to detect coevolving sites [33] does not identify contacting residues in evolutionary trajectories simulated by RosettaEvolve (data not shown). The evolutionary basis for sequence covariation is therefore not fully understood. Talavera et al. [30] argued that coordinated sequence changes require very high selective pressures to occur, which results in rates so slow that coevolution would not be measurable. They argue that covariation is the consequence of sites with slow evolutionary rates rather than coevolution. Given the practical importance of statistical coupling methods in bioinformatics, it is of great interest to understand the relationship between covariation, coevolution, and protein energetics.

In this study, we investigate whether covariation signals emerge in sequences simulated from a phylogenetic tree using a detailed atomistic simulation of protein energetics and how the strength of the selection pressure affects the covariation signal. To address these questions, we inferred a phylogenetic tree from an alignment of the natural sequence of azurin and used it as the basis for evolutionary simulations with RosettaEvolve. Simulated phylogenetic trees were generated at different selection pressures, starting from equilibrated sequences at each given selection pressure. We developed a recursive algorithm that generates evolutionary trajectories over a given tree topology and branch lengths. We populated the tree 11 different times with variable selection pressure.

The sequence entropy at the leaves of simulated azurin trees depends strongly on the selection pressure (Fig. 4). Using parameter values we previously found to explain natural amino acid substitution patterns [5, 11], we see similar position-specific sequence entropies between our simulated proteins and their natural counterparts.

**Figure 4:**
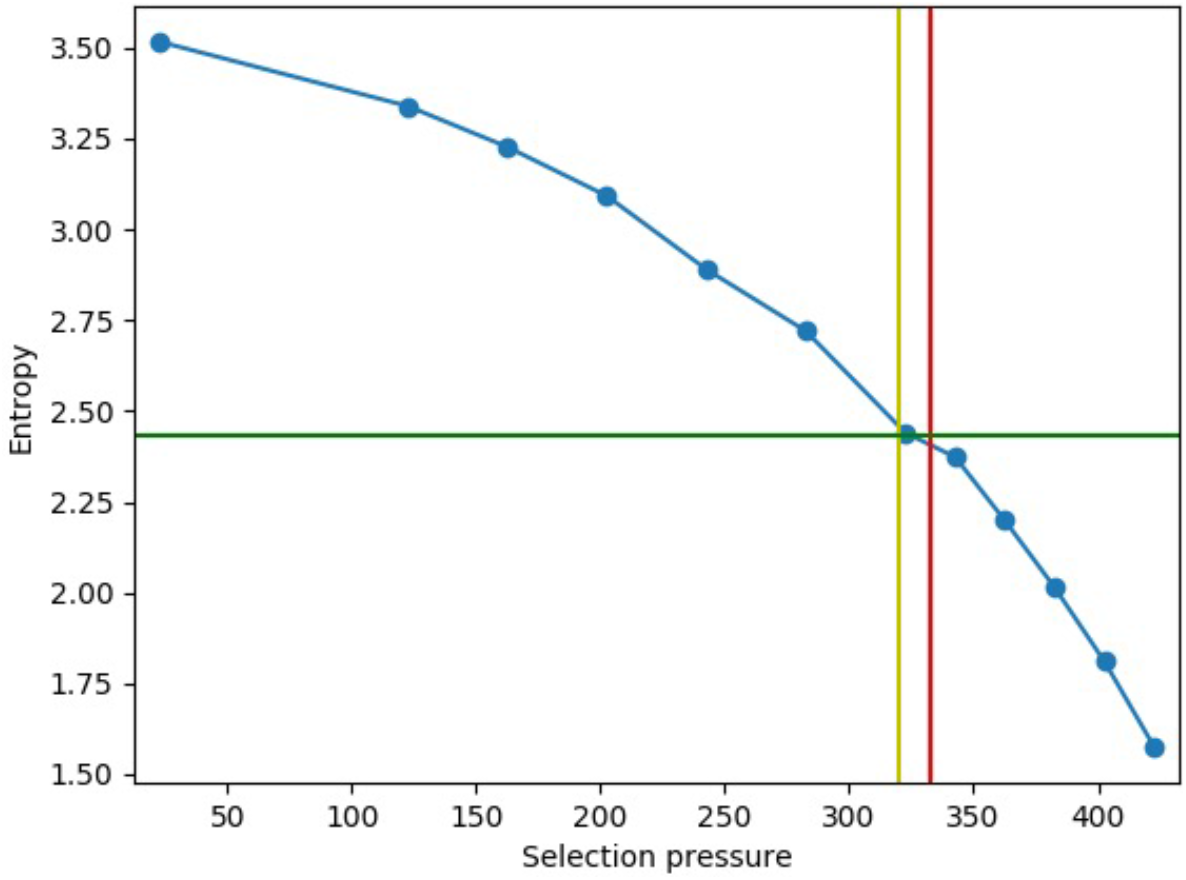
Simulation of phylogenetic trees of azurin with RosettaEvolve. Dependence of sequence entropy of leaf sequences with reference energies. The green line corresponds to the entropy in the natural sequences used to infer the azurin tree. The red line corresponds to the energy of the native sequence of azurin. The yellow line corresponds to the selection pressure that maximizes the correlation between computed and empirical amino acid substitution rates in Norn et al.[7].

The leaf sequences generated by RosettaEvolve trajectories simulated at different selection pressure values were analyzed for covariation signal statistical coupling score using Gremlin [34]. The ability of Gremlin in predicting residue-residue contacts was summarized in Receiver Operator Characteristic (ROC) curves, where the true positive rate is plotted against false positive rate. In Figure 5, the ROC curve for the natural sequences is compared to sequences simulated at two different selection pressures, one corresponding to low (red curve) and one to high selection pressure (orange curve). The area under the curve (AUC) is a metric for the overall performance. For the high selection pressure simulations, the AUC reaches the same values as the natural sequences, while sequences evolved with low selection pressure provide a considerably worse basis for predicting residue-residue contacts.

**Figure 5:**
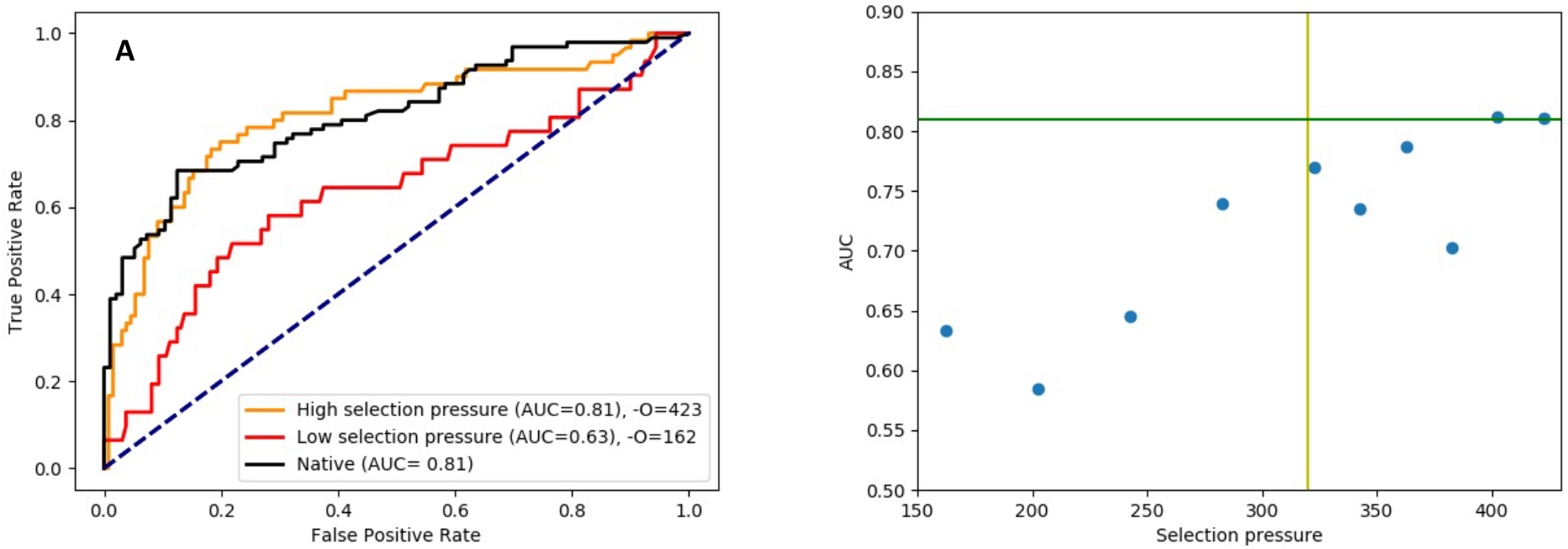
ROC curve for residue-residue contact prediction. A) Comparison of ROC curve for natural sequences (black) and two simulated alignments (red and orange) at two different reference energies. B) Dependence of contact-prediction accuracy (AUC) on selection pressure. The green line corresponds to AUC for the natural sequence. The yellow line corresponds to the selection pressure that gives the optimal correspondence between predicted and empirical amino acid substitution rates in a Rosetta-based rate prediction method [7, 16]. The yellow line corresponds to the selection pressure that maximizes the correlation between computed and empirical amino acid substitution rates in Norn et al.[7]. The blue line shows the diagonal.

Even though similar AUC values are found for some simulated sequences, the early enrichment is nonetheless better for the natural sequence, resulting in higher prediction accuracy in the range relevant for structure prediction. The overall predictive power (characterized by AUC) is highly dependent on the selection pressure. In Figure 5B, the AUC is plotted against selection pressure. The ability of GREMLIN to identify true residue-residue contacts drops with decreasing protein stability (as controlled by the selection pressure). For reference stabilities corresponding to most stable proteins, up to 78% (43 out of 55 contacts above the threshold used by Gremlin to predict contacts) of the predicted residue-contacts contacts are validated in the structure corresponding to the native sequence of azurin.

We calculated residue-residue pair energies from the crystal structure of azurin using Rosetta, see Figure 6. The average pair energy between residues predicted to be in contact by Gremlin (blue distribution) has pair interaction values that are considerably stronger than contacts in general in the protein (grey distribution). The mean interaction energy is around −1 kcal/mol for sequences simulated with stabilizing reference energies for the predicted Gremlin contacts, compared to −0.15 kcal/mol for all contacts in the protein. So, for the most stable proteins, contacts detected by statistical coupling analysis correspond to pair interactions among the most stabilizing contacts in the protein.

**Figure 6:**
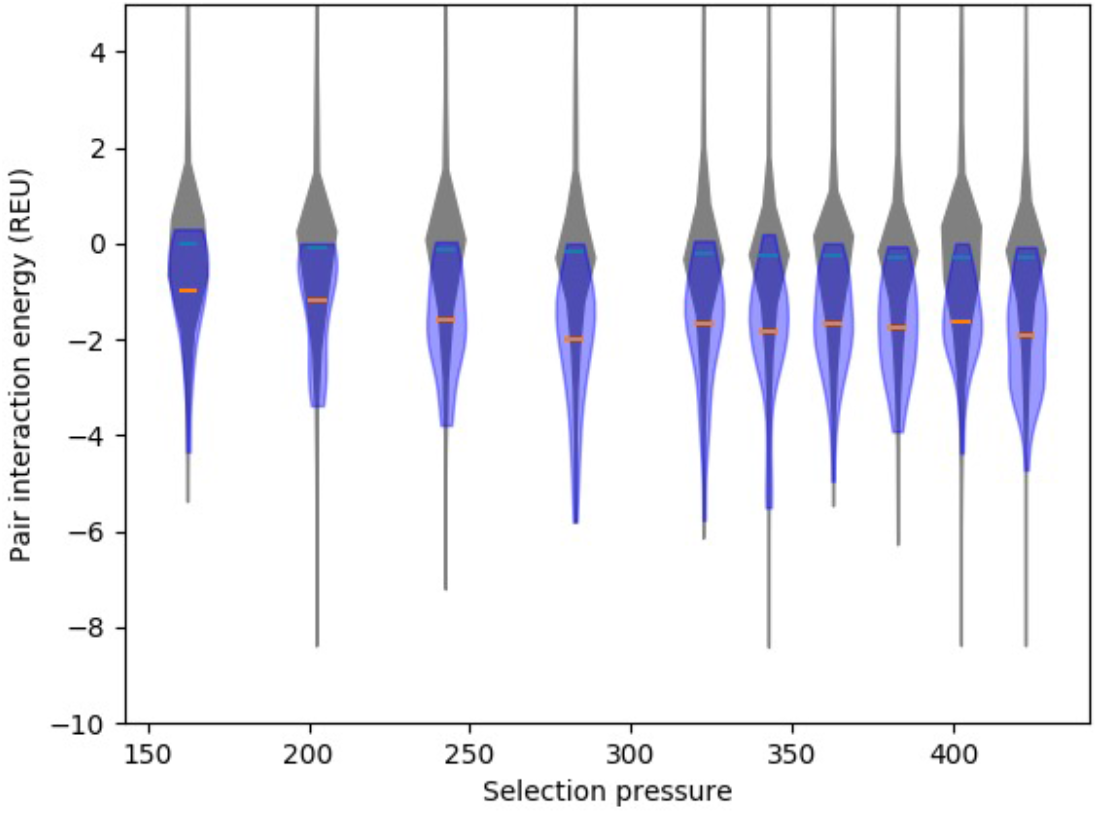
Pair interaction energy for contacts predicted from alignments. Distribution of pair interaction energies for contacts predicted by Gremlin (blue) (orange lines mean) and all contacts within the structure (gray, green line) as a function of selection pressure. The width of the violin is related to the frequency of a given pair interaction value. Energies in Rosetta Energy Units (REU).

We further analyzed the evolutionary trajectory of contacts detected by the statistical coupling analysis. Starting from the leaf nodes, we identify the branch point where a residue pair found in the leaf node was first introduced and characterize the change in energy on the evolutionary path towards the leaf node. We find that the average change in energy for the two residues in the predicted contact (in the context of the entire structure) is favorable, but only slightly so (−0.15 kcal/mol). Thus, when the pair was formed, there did not appear to be a large energetic gain in forming the contact. However, the selected pair may become energetically entrenched after initially appearing (evolutionary Stokes shift), or there may be special conditions before it was inserted. Further analysis of the fluctuation in selection coefficients over time will have to be carried out to fully understand the mechanism behind covariation signals for these residue pairs.

#### Fluctuations in protein stability result in fluctuations in site rates

During the evolutionary trajectories, sites in the protein will experience a fluctuating structural environment. How much do site rates fluctuate during a mutational trajectory? How much of this variation can be explained by fluctuations in protein stability during an evolutionary trajectory? We calculated site-specific rates across an evolutionary trajectory corresponding to a branch length of 1 mutation per site to address these questions. After each mutation, we calculated the energy, site-rates and compared site rates to empirical values predicted by rate4site [35] from the sequence alignment of azurin. Figure 7A shows the energy and correlation with empirical site rates fluctuate across the trajectory. The Pearson correlation between calculated and empirical site-specific rates fluctuates considerably during the trajectory, ranging from 0.48 to 0.61. Fluctuations in the stability of the protein (Figure 6A, red line) will result in an overall change in substitution rate, with less stable proteins having higher mutation rates. About 31% of the variation in the correlation with empirical site rates can be explained by the fluctuation in stability of the protein during the trajectory.

**Figure 7:**
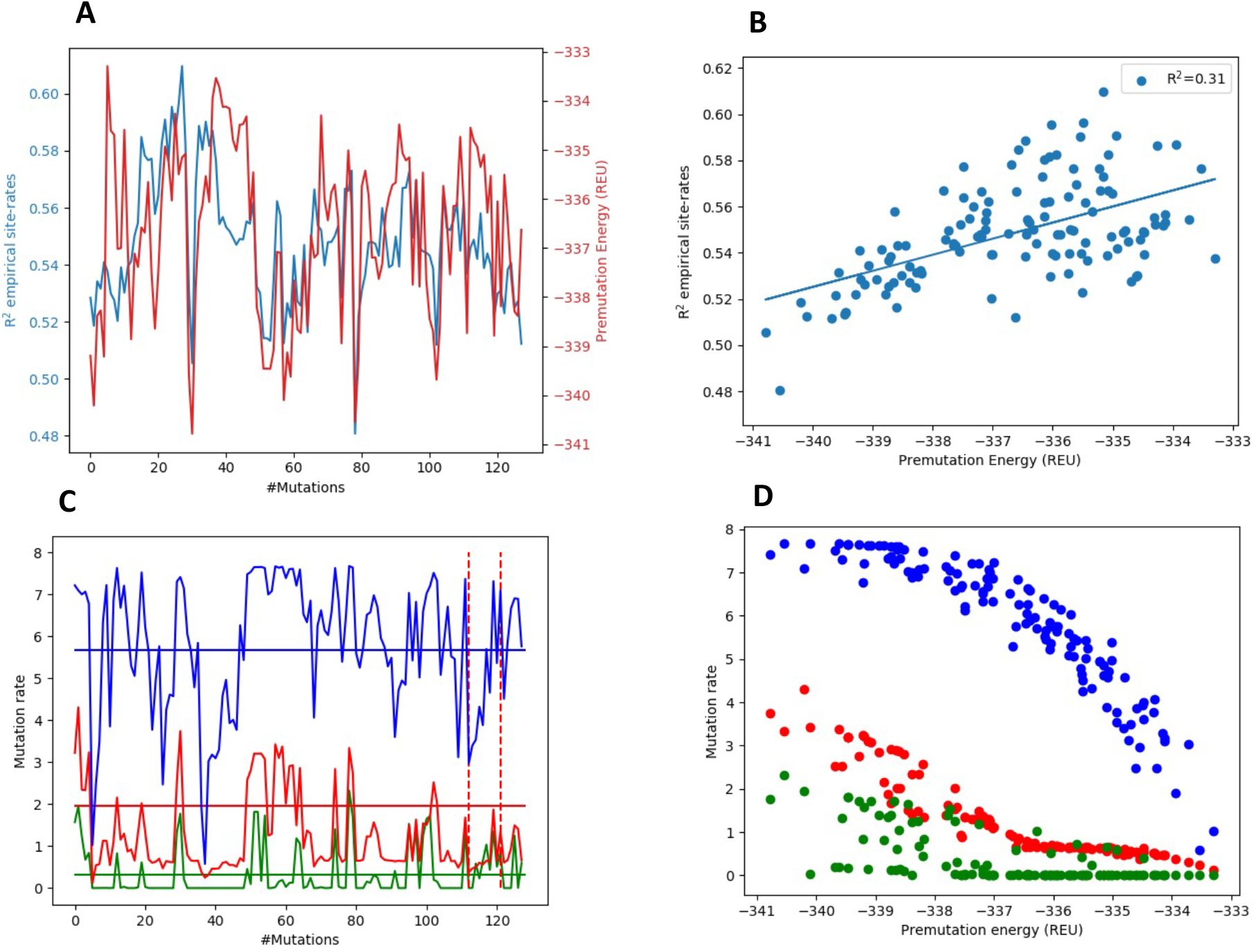
Fluctuation in mutation rates during a trajectory and correlations with stability. A) Correlation between calculated and empirical site-specific rates (calculated as R^2^-values) for azurin (blue) as a function of the number of introduced mutations. Fluctuation in energy as a function of introduced mutations (blue). B) Correlation between R2-values and energies are shown in A). Selection pressure in trajectory is set to −322. C) Site-specific rates for the sites in azurin. Residue 41 (blue), residue 49 (green) and residue 66 (red). Empirical site rates from rate4site as solid lines and dashed lines show mutational events at the site. D) Dependence of site-rates in C) with premutation stability. Energies in Rosetta Energy Units (REU).

The rate at individual sites in the protein will also fluctuate considerably. In Figure 7C, the rates for three different individual sites in the protein are plotted as a function of the number of mutations in the trajectory. The mutation rate can drop an order of magnitude during the trajectory, even though there are no mutations occurring at that site. Figure 7D demonstrates that rates at individual sites can be highly coupled to the stability of the proteins.

## Discussion

Protein stability results from the net balance of forces involving thousands of interacting atoms. The result is typically a protein with only marginal stability where small changes in atomic interactions can shift the protein from a folded to unfolded state. The marginal stability of proteins can be understood as the result of the balance between the introduction of predominantly destabilizing mutations and selection [14]. This marginal stability also emerges in evolutionary simulations employing a simple contact-based potential of protein energetics [14]. Nonetheless, many mechanistically important aspects of protein evolution may be lost without consideration of the detailed atomic interactions in proteins.

A few investigations have been presented where evolutionary trajectories have been simulated with atomistic energy functions. Typically, these studies have employed the FoldX energy function[13] to evaluate DDG values [14, 15]. In this manuscript, we simulate evolution with the Rosetta macromolecular modeling package, which provide a powerful framework for modeling structure and energetics of proteins[16] and where side-chain and backbone flexibility can be modeled with a wide range of structure-prediction protocols. RosettaEvolve can be readily extended to use additional fitness models and additional methods to model conformational changes upon mutations, such as the flexible backbone approach developed by Bartlow et al. to model mutations in protein interfaces[17].

Jiang et al. [18] used Rosetta to generate evolutionary trajectories along a single branch. There are several differences between their approach and ours. We simulate evolution based on nucleotide mutations, rather than at the protein level, we use a DDG calculation that includes more backbone flexibility, and our models differ in the selection pressure and effective population size. Jiang et al. [18] used protein design calculations to set the selection pressure value and used a smaller effective population size (100).

We have previously developed a Rosetta-based method to predict amino acid substitution rates[5] from protein structure using the combination of structure-based stability calculations and mutation-selection model, which we refer to as the TMS (Thermodynamic Mutation-Selection) model. Amino acid substitution rates at a site calculated can readily be summed up to evaluate the site substitution rates[18]. A benefit of the TMS method is that it enables us to evaluate the site-specific rates for all sequences continuously along an evolutionary trajectory. Our results show that the site rates fluctuate considerably during the trajectory, even for sites that are not mutated. Natural proteins have also experienced significant variation in backbone structure during their evolutionary trajectories as reflected by the structural variability found in sequence homologs. Such relatively large-scale structural fluctuations are not modeled with the limited backbone flexibility DDG method used in this study. Our simulation results highlight that relatively small changes in structure and energetics in proteins can have considerable consequences for substitution rates at individual sites in proteins and that accurate prediction of site-rates hinges on modeling the detailed structural and energetic consequences of amino acid substitutions. Nonetheless, a significant amount of the fluctuation in substitution rates can be explained simply by fluctuations in the overall stability of proteins during the evolutionary trajectory (Fig. 7BD).

We show that phylogenetic trees populated with sequences using an evolutionary all-atom structural and energetic model result in sequences with a significant covariation signal. Sites with high statistical coupling have considerably more favorable pair interaction energies than average contacts in proteins. This suggests that the basic premise behind statistical coupling analysis for contact prediction - that strong residue-residue interactions lead to covariation signal - is correct. Nonetheless, although some covariation signal is also observed at lower selection pressures, only at very high fitness pressures does the covariation signal reach the levels seen for natural sequences. Furthermore, the limited backbone flexibility in the simulation likely overestimates the relative strength of specific residue-residue interactions, resulting in enhanced covariation signals. We, therefore, expect that more realistic modeling of structural variability would reduce the covariation signal. Taken together, this may suggest that additional mechanisms can be behind the strong covariation signal found in natural protein sequences. Further investigations of the correlation between substitution history of RosettaEvolve trajectories, statistical coupling score, and protein energetics should enable a more detailed understanding of how covariation emerges among homologous proteins.

## Materials and Methods

### Structural modeling

The crystal structure of azurin (PDB ID: 5AZU) was used as the basis for all modeling. All structural modeling was done with the Rosetta macromolecular modeling suite[16] using the beta_nov16 energy function. 5AZU was energy refined before running the evolutionary trajectories using the method described by Niven et al.[19] to make the crystal structure compatible with the energy function. Prediction of DDG values for mutations was done using a modified version of the approach presented by Park et al. [10], with a 6.0 instead of 9.0 Å distance cutoff in the Lennard-Jones energy term. Backbone flexibility is allowed at the mutated and neighboring residues, and side-chains are repacked for all residues that have at least an interaction energy more than 0.1 kcal/mol.

### Site-specific rate calculations

Site rates were calculated with the TMS method presented in Norn et al. [7] as described in [41]. In reference [41] a single selection pressure was fitted for a benchmark of 66 proteins based on maximizing the similarity with empirical site-specific rates. In this study, the selection pressure used in the rate calculation corresponds to the value used in the evolutionary trajectory that generated the structure. Empirical rates for azurin were calculated with rate4site [35] using the empirical Bayes method with the LG instantaneous rate matrix and an alignment consisting of 500 sequences.

### Tree inference phylogenetic tree simulation

A phylogenetic tree was generated based on a sequence alignment generated by Gremlin [20] using RAxML[21] with the LG as the instantaneous rate matrix.

Before running the tree crawling algorithm the effective mutation rate of RosettaEvolve for a given selection pressure was determined by running trajectories with both synonymous and non-synonymous mutations enabled. By measuring the number of non-synonymous mutations per mutational trials in the simulation the mutation rates were calculated as

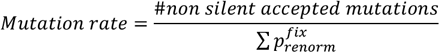

where 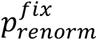 corresponds to the amount of time before fixation at each step in the trajectory.

Filling of the phylogenetic tree was done from a node at the center of the tree to optimize computational speed. For each branch RosettaEvolve was run with the number of mutations expected from the branch length in the empirical tree. At each internal node a structure is stored and use as a basis for the next set of branches originating from each leaf.

### Statistical coupling analysis

Leaf sequences from the phylogenetic tree simulation (1050 sequences) were analyzed with Gremlin[20] web server (gremlin.bakerlab.org). Gremlin was run without MSA enrichment so that only simulated sequences was used in the analysis. Classification of contact prediction was done using the standard distance threshold of 8.0 Å between Cb (Ca for glycine) using the coordinates in 5AZU. A default threshold value of a scaled score above 1.0 was used to select contacts predicted by Gremlin.

Pair energies were determined using the residue_energy_breakdown.linuxgccrelease application using the beta_nov16 energy function and the energy-refined 5AZU structure.

### Command lines and code

RosettaEvolve will be made available through a release of Rosetta[16]. Additional scripts and running information can be found at https://github.com/Andre-lab/RosettaEvolve. Command lines used in this study are found in Supplementary materials.

## Acknowledgements

We thank Douglas L. Theobald for helpful discussions on the implementation of RosettaEvolve. This work was funded by a grant from the Swedish research council (grant number 2015-04203) to IA. The computations were enabled by resources provided by the Swedish National Infrastructure for Computing (SNIC), partially funded by the Swedish Research Council through grant agreement no. 2020:5-308.

## Notes

### Competing Interest Statement

The authors have declared no competing interest.

